# NF-κB c-REL-OTUD4 axis regulates B-cell receptor in B-cell lymphoma

**DOI:** 10.1101/2023.05.06.539691

**Authors:** Eslam Katab, Anushree Jai Kumar, Katja Steiger, Julia Mergner, Mikel Azkargorta, Assa Yeroslaviz, Felix Elortza, Vanesa Fernández-Sáiz

## Abstract

The B-cell receptor (BCR) is essential for B-cell development and a crucial clinical target in immuno-oncology. However, therapeutic success against the BCR and downstream signaling pathways is hampered by enhanced NF-κB activation as a resistance mechanism. Using a multiomic approach, we discover the c-REL proto-oncogenic subunit of the NF-κB family as a key transcription factor regulating BCR subunit levels in B-cell lymphoma. Subsequent ChIP-seq, cell biology experiments, and patient data analysis reveal that OTUD4 is a critical deubiquitinase for inhibiting proteasomal degradation of c-REL and for stabilizing a multi-loop positive feedback of NF-κB to the BCR pathway. Remarkably, *OTUD4* downregulation destabilizes c-REL and BCR levels and inhibits cell growth of B cell lymphoma. Thus, we shed light on the malignant potential of c-REL abundance, identify a positive feedback from c-REL to upstream BCR and present OTUD4 as a vulnerability to synergistically target NF-κB and BCR pathways in B-cell lymphoid malignancies.

## Introduction

A unique property of B-lymphocytes is the presence of the B-cell receptor (BCR) on the cell surface consisting of an extracellular antigen recognizing domain (a transmembrane antibody) and two intracellular signal transducing subunits (CD79A and CD79B)^1, 2^. Signaling pathways emanating from the BCR regulate most functions of B cells and are essential for their development and maturation^3–5^. In lymphoid malignancies, constitutive B-cell receptor activities are frequently observed to promote aberrant proliferation and survival of B-cells^6–8^. Thus, numerous clinical therapies are directed to target BCR and their downstream effectors^9, 10^. However, the development of resistant mechanisms such as NF-κB gene signatures crucial for cell survival, immune evasion and epigenomic reprogramming, challenge clinical therapies in an unclear manner^11–13^. Under unstimulated conditions, NF-κB inhibitory proteins (IκBs) sequester NF-κB transcription factors in the cytoplasm. On the contrary, upon multiple activation signals including stimulation of receptors on the plasma membrane, the ubiquitin proteasome system (UPS) critically regulates NF-κB activation by proteasomal degradation of the IκBs^5, 14, 15^. The release of NF-κB subunits allows transport of NF-κB dimers into the nucleus to regulate transcription of target genes. However, cumulative evidence shows that NF-κB transcription factors are subjected to additional regulation beyond that exerted by IκBs^16, 17^. Of notice, while signal transduction from membrane receptors to activation of the NF-κB pathway in the nucleus has been extensively reported, a limited number of studies have described NF-κB-mediated control of gene expression to impact at cell surface receptors, consisting in many cases of negative feedback through the transcription of "decoy receptors" unable to relay signals to downstream targets^18^.

The proto-oncogene c-REL is a unique member of the NF-κB transcription factor family with capacity to malignantly transform chicken lymphoid cells and to regulate the immune response of myeloid cells and lymphocytes^19–24^. The oncogenic potential of c-REL is determined by its C-terminal transactivation domain (TDA) that shows little similarity with other members of the NF-κB family and seems to be responsible for each proteińs unique function^25–27^. Unlike other NF-κBs expressed in many cell types, expression of c-REL is restricted to hematopoietic and epidermal cell lineages, being enhanced in B and T cells. The prominent role of c-REL in lymphoma and autoimmune disease is underscored by its frequent activation as well as amplification and gain of the *REL* gene locus^28–31^. Accordingly, considerable effort has been made to develop strategies to inhibit NF-κB members including c- REL^21, 32–34^. Therefore, understanding on the one hand which genes are specifically regulated with the intervention of the TDA of c-REL, and on the other hand the molecular mechanisms that govern *c-REL* expression and c-REL abundance, are of high interest in immuno-oncology. In regard to c-REL protein abundance, ubiquitination at the C-terminal region of v-Rel, the viral oncogenic version of c-REL, was described two decades ago to regulate its proteasomal degradation in T cells^35^. However, the C-terminal lysine residues of v-Rel are not conserved in human homologous. More recently, proteasomal inhibition was described to stabilize ubiquitinated c-Rel in macrophages^36^ and the E3 ubiquitin ligase Peli1 was reported to mediate proteasomal degradation and inhibition of c-Rel nuclear translocation in T cells^37^. Nonetheless, the molecular mechanism(s) to mediate ubiquitin-mediated degradation of c-REL in B-cell lymphomas, as well and their biological consequences, have not been explored.

We reveal NF-κB c-REL as an essential transcription factor to regulate the abundance of BCR receptor subunits in B-cell lymphoma. Fine-tune regulation of c-REL levels is achieved by deubiquitinase *OTU domain-containing protein 4* (OTUD4) that stabilizes c-REL in a feedback loop mechanism. Whereas c-REL modulates the transcriptional regulation of *OTUD4*, the deubiquitinating activity of OTUD4 counteracts protein turnover of c-REL and allows for c-REL nuclear accumulation. By identifying the c-REL-OTUD4 axis to stabilize the BCR in B-cell lymphoma, we discover a positive multi-loop feedback from NF-κB to upstream BCR and reveal OTUD4 as a vulnerability to synergistically target NF-κB and BCR pathways in B-cell malignancies.

## Results

### Multi-omics reveal *c-REL* to regulate BCR subunits in B-cell lymphoma

To better understand the role of proto-oncogenic c-REL in lymphoid cancer cells, we carried out a multiomic approach combining *c-REL*-dependent transcriptome and whole proteome analysis, that could lead to non-redundant readouts^38, 39^. For our initial analysis, we employed the phenotypically germinal center B-cell like DLBCL (GC-DLBCL) Burkitt lymphoma BJAB cell line, where c-REL over-expression was described to enhance their transformation state^40^. Validated control-shRNA or up to three different *c-REL*-targeted shRNAs were employed to down-regulate *c-REL*. BJAB cells were analyzed in total RNA sequencing (RNA-seq) or whole Mass-Spectrometry (MS) quantitative experiments. Among the genes and proteins of relevance in DLBCL that showed a significant variation upon *c-REL* downregulation, we found Immunoglobulin heavy constant mu (IGHM), the heavy chain of the BCR in BJAB cells^41^, to be substantially downregulated both at the mRNA and protein levels (Figs. 1A-B; Supplementary Figure 1; Supplementary Tables 1 and 2). A significant decrease of the much smaller CD79B cytosolic subunit of the BCR was also found by MS (Fig. 1A). To discard aberrant immunoblotting of large immunoglobulin transmembrane protein domains^42^, we confirmed our results by WB of the intracellular CD79B subunit (Figs. 1C and 1D). In a complementary over-expression experiment, CD79B levels were substantially increased upon overexpression of c-REL (Figs. 1E and 1F). Intrigued by these findings in BJAB cells, we corroborated by WB the decrease of CD79B upon *c-REL* downregulation also in HBL-1 and OCI-Ly3 DLBCL cell lines (Supplementary Figs. 2A-B). Our results show a role of the NF-κB transcription factor c-REL in the regulation of BCR subunits in B-cell lymphoma cells.

**Fig. 1.**
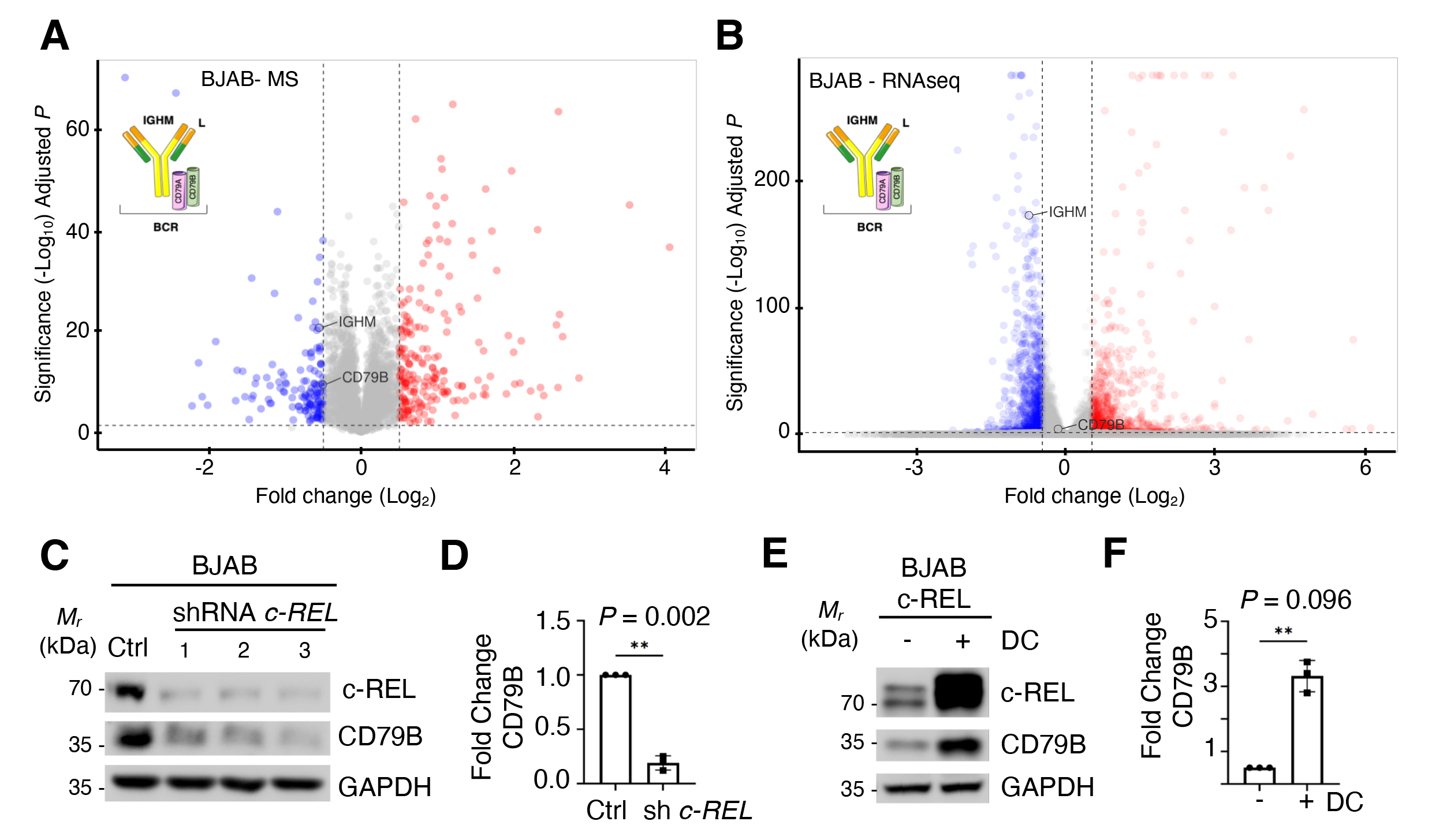
Multiomic quantitative analysis reveals *c-REL* as a regulator of BCR levels. **A)** Proteomic analysis. Volcano plot of proteins significantly downregulated (blue), upregulated (red) or with no significative difference (grey) in the total proteome samples of BJAB cells upon *c-REL* downregulation *vs* control cells measured by quantitative-MS. Each dot represents one protein. Significance threshold is set at 1.3 (-Log_10_) and fold change threshold is set between −0.5 and +0.5 (log_2_). Dots corresponding to the IGHM and CD79B subunits of the BCR are depicted. Biological triplicates using two different shRNAs against *c-REL* were used in the measurements. A scheme of the BCR complex is included. See Supplementary Table 1. **B)** Transcriptomic analysis. Volcano plot of mRNAs of genes significantly downregulated (blue), upregulated (red) or with no significative difference (grey) in total RNA samples of BJAB cells upon *c-REL* downregulation vs control cells measured in RNA-seq. Each dot represents one gene. Significance threshold is set at 1.3 (-Log_10_) and fold change threshold is set between −0.5 and +0.5 (log_2_). Dots corresponding to the *IGHM* and *CD79B* subunits of the BCR are depicted. Biological triplicates using two different shRNAs against c-REL were used. A scheme of the BCR complex is included. See Supplementary Table 2. **C)** Western blot analysis of BJAB cells upon scrambled shRNA control or *c-REL* targeted shRNAs. c-REL, CD79B, or loading control GAPDH proteins are blotted against corresponding antibodies. **D)** Quantification of CD79B decrease in *c-REL* downregulated cells *vs* control cells shown in C. The error bar represents the mean ± S.D, (n = 3). Significance is indicated. **E)** Western blot analysis of BJAB cells upon c-REL over-expression using an inducible expression system in absence (-) or presence (+) of doxycycline (DC) inducer for 24h (1ng/ml). c-REL, CD79B, or GAPDH proteins are blotted against the indicated antibodies. **F)** Quantification of CD79B increase in cells over-expressing *c-REL* vs control cells shown in E. The error bar represents the mean ± S.D, (n = 3). Significance is indicated.

### c-REL regulates OTUD4 expression and correlates with OTUD4 in DLBCL

Prompted by the previous findings, we aimed to dissect the direct target genes under c-REL transcriptional control. To that end, we first investigated the c-REL binding sites at the genomic level by chromatin immunoprecipitation followed by sequencing (ChIP-seq) in BJAB cells. Using a c-REL antibody previously validated against the more specific C-terminal region of c- REL^43, 44^ (Fig. 2B), we found *OTUD4* deubiquitinase as a prominent hit (Fig. 2A; Supplementary Table 3). Next, we performed qPCR experiments that confirmed the presence of *OTUD4* (Fig. 2C). Consistently, OTUD4 protein levels were increased under c-REL overexpression (Supplementary Fig. 3). Noticeably, among the approximately 100 deubiquitinases (DUBs) present in the human genome, *OTUD4* mRNA showed the most correlative result with *c-REL* in hematologic cells in the GEPIA database^45^ (Fig. 2D; Supplementary Table 4). Of further importance, among hematologic malignancies *OTUD4* appears upregulated in DLBCL^46^ and it is associated with a poorer prognosis (Supplementary Figs. 4A-C). To corroborate the *in-silico* data we performed immunohistochemical staining of OTUD4 in a patient cohort (n = 55) of DLBCL and analyzed its correlation with c-REL- staining. Remarkably, we found a positive correlation of OTUD4 and c-REL in DLBCL patient immunostaining (Figs. 2E-F; Supplementary Fig. 4D).

**Fig. 2.**
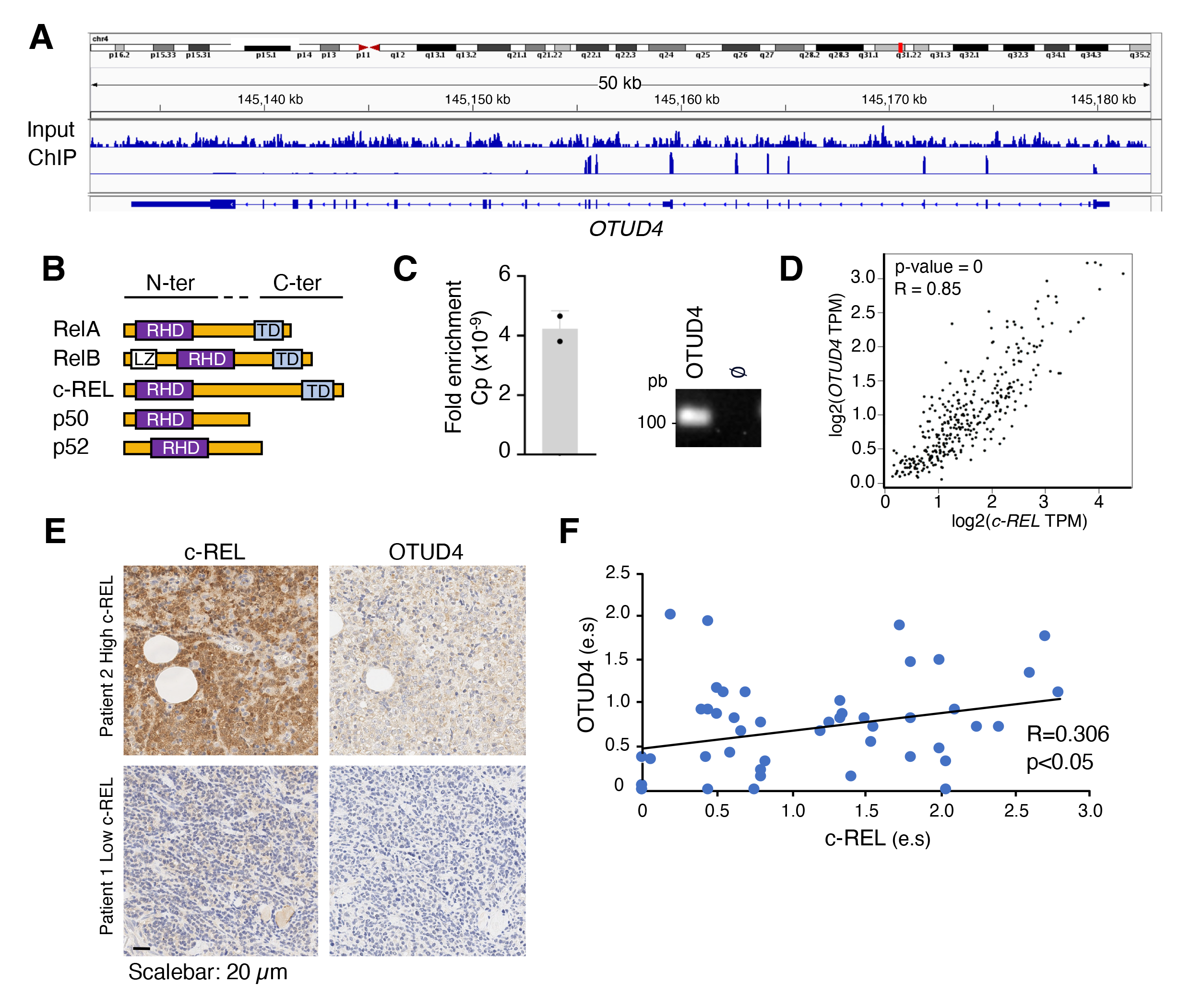
*OTUD4* is a target of c-REL that correlates with c-REL in DLBCL. **A)** ChIP-seq data showing c-REL binding in the region of the *OTUD4* gene in BJAB cells. See Supplementary Table 3 for a detailed information of peaks. **B)** Schematic representation of NF-κB subunits in humans. Conserved Rel Homology domain (RHD) and Leucine zipper (LZ) in the N-terminal regions, or the divergent Transactivation domains (TD) in the C-terminal region of proteins are depicted **C)** qPCR enrichment (link) of *OTUD4* gene visualized by representative gel-electrophoresis (right) in ChIP samples in Fig. A. The error bar represents the mean ± S.D, (n = 3). **D)** Correlation of *OTUD4* mRNA levels with *c-REL* in whole blood cells in GEPIA database. Normalized counts in "Transcripts per kilobase per million" (TPM) of *OTUD4* and *c-REL* to correct for transcript length and library size are shown. Significance is indicated. See Supplementary Table 4. **E)** Representative histological analysis of DLBCL patient tissues (n= 55) stained with c-REL and OTUD4 antibodies. **F)** Correlation analysis between the expression score (e.s) of OTUD4 and c-REL signals. Analysis was done by SPSS 28.0 software. See Supplementary Fig. 4 for details.

Together, our data show that c-REL mediates transcriptional control of *OTUD4* in B-cell lymphoma and correlates with OTUD4 in DLBCL.

### OTUD4 counteracts proteasomal degradation of c-REL and mediates its nuclear localization

OTUD4 was previously found to interact with c-REL in a proteome-scale map and to regulate NF-κB activity in Mouse Embryonic Fibroblasts (MEFs)^47, 48^. We therefore tested the hypothesis that OTUD4 might interact c-REL also at the protein level and function as a DUB for c-REL. Because both c-REL and OTUD4 harbor nuclear localization signals (Fig. 3A), and have been described to translocate between the cytoplasm and the nucleus^24, 48, 49^, we first analyzed the sub-localization of OTUD4 and c-REL in B-cell lymphoma fractionated cell extracts. Endogenous OTUD4 levels in DLBCL cells showed a predominant cytoplasmic localization while the localization of c-REL was dependent on the B-cell lymphoma subtype, in good accordance with previous data^50^ (Fig 3B). We first validated the OTUD4 interaction to c-REL by immunoprecipitation experiments using overexpressed proteins in HEK293T cells. Samples of immunoprecipitated OTUD4, but not OTUD6B as a control DUB of the OTU domain-containing family, specifically interacted with c-REL (Fig. 3C). We then analyzed the OTUD4 interaction with c-REL in fractionated cell extracts. In accordance with previous findings, overexpression of c-REL leads to its nuclear accumulation^51, 52^ while most of the overexpressed OTUD4 appeared in the cytoplasm^53^ (Fig. 3D). We detected a clear interaction of OTUD4 and c-REL in the cytoplasmic fraction (Fig. 3D). In following experiments, we tested the ubiquitination status of c-REL in the cytoplasmic and nuclear fractions of activated B-cell (ABC) HBL-1 and TMD8 DLBCL cell lines that show a comparable amount of c-REL levels in both cellular compartments. Using GST-TUBEs, we detected a c-REL signal mostly in cytosolic extracts compared to nuclear fractions of DLBCL cells (Figs. 3E-F). Together, these experiments suggest that c-REL interaction to OTUD4 occurs mainly in the cytosolic fraction, where ubiquitin is associated to c-REL.

**Fig. 3.**
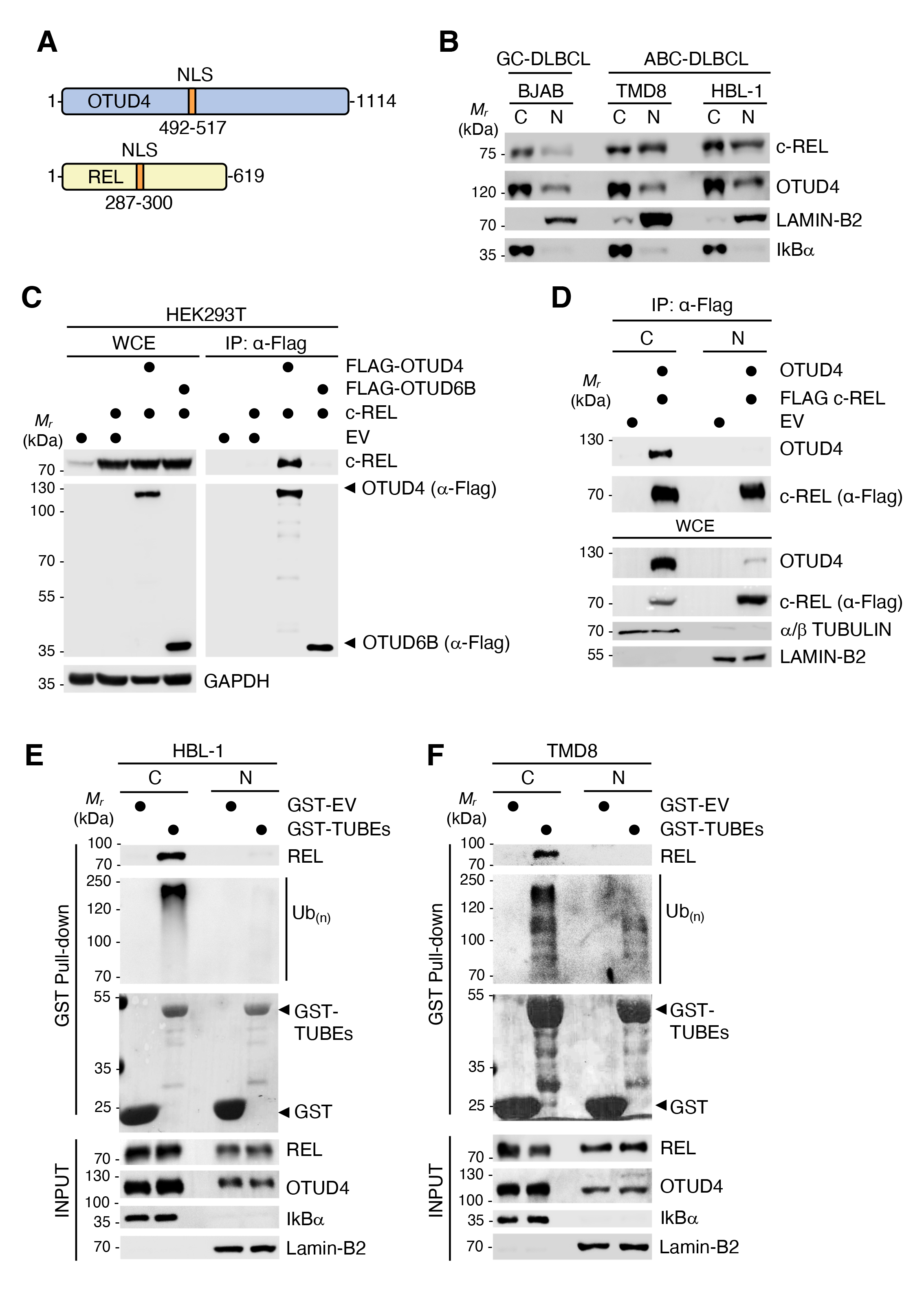
c-REL is ubiquitinated in DLBCL and interacts with OTUD4. **A)** Scheme of OTUD4 and c-REL proteins. Nuclear localization signals (NLS) according to Myhits server (https://myhits.sib.swiss) are indicated in orange. **B)** Sublocalization of endogenous c-REL and OTUD4 in DLBCL cell lines. Fractionation of cytoplasmic (C) and nuclear (N) extracts of GC-DLBCL BJAB, or ABC-type TMD8 and HBL-1 cell lines. Endogenous levels of c-REL, OTUD4 or the nuclear LAMIN-B2 and cytoplasmic marker IκBalpha were blotted against the corresponding antibodies. **C)** OTUD4 interaction with c-REL. Western blot of whole cell extracts (WCE) and FLAG-beads immunoprecipitations (IP) from HEK293T cells transiently overexpressing FLAG-tagged OTUD4 or FLAG-OTUD6B as a control. **D)** Subcellular interaction of OTUD4 with c-REL. Cell lysates of HEK293T with overexpressed FLAG-tagged c-REL and non-tagged OTUD4, as indicated, were fractionated into cytoplasmic (C) and nuclear (N) fractions. FLAG-c-REL immunoprecipitation was performed using anti-FLAG beads. Captured proteins were blotted against the depicted antibodies. **E)** Pull downs of ubiquitinated proteins in DLBCL HBL-1 cells. Cytoplasmic or nuclear extracts of DLBCL HBL-1 cells were pulled-down with GST or GST-TUBEs coupled to glutathion beads. Proteins bound to beads were blotted against ubiquitin and c-REL antibodies. **F)** Same experiment as in E performed with DLBCL TMD8 cell extracts.

To confirm that OTUD4 mediates ubiquitin cleavage from c-REL, we performed c-REL de-ubiquitination assays in HEK293T cells upon c-REL and OTUD4 overexpression. We observed a clear decrease in the total ubiquitination signal of c-REL in OTUD4 overexpressing cells *vs* control cells (Figs. 4A, B). Upon *OTUD4* downregulation, on the contrary, the total ubiquitin signal associated to REL was increased (Figs. 4C, D). To further exclude that the ubiquitination signal observed in our IPs experiments under denaturing conditions originates from a c-REL interactor, we created a BJAB cell line expressing HIS_(6x)_-tagged-ubiquitin and silenced *OTUD4*. Pull down-experiments of ubiquitinated proteins under stringent denaturing conditions showed the appearance of a higher molecular weight band of c-REL bound to the nickel beads that was enhanced in the case of *OTUD4* downregulation compared to control cells (Figs. 4E- G). Of notice, a higher molecular weight band of c-REL was diminished in the total cell extracts of *OTUD4*-silenced cells (Fig. 4F). This result suggested that *OTUD4* downregulation decreases the levels in the total cell lysate of a post-translationally modified form of c-REL with higher affinity for nickel beads, likely due to rapid degradation by the proteasome. To validate this hypothesis, we performed a time-course stability assay of c-REL in the presence of cycloheximide under downregulation of *OTUD4* and proteasomal inhibition as indicated (Fig. 4H). In the scrambled shRNA control cells, we detected a clear time-dependent decrease of the higher molecular band of c-REL. Importantly, this higher molecular band of c-REL was rescued with proteasomal inhibition (Figs. 4H, 4I). In contrast, we observed a lower band corresponding to the expected size of c-REL that remained stable in the course of the experiment. In good accordance with the role of OTUD4 as a deubiquitinating enzyme for c-REL, the higher MW band of c-REL rescued by MG132 treatment was absent in *OTUD4* downregulated cells, whereas the stable lower MW band was diminished (Fig. 4H). Loss of *OTUD4* did not show any effect on the stability of MYD88, a described substrate of the OTUD4 activity on ubiquitin linkage-K63^48^, while a decrease of CD79B levels was observed (Fig. 4H). To further exclude the possibility that the effects observed in c-REL were due to indirect modulation of OTUD4 via MYD88 signaling, we inhibited MYD88 in a control experiment that did not show variation on c-REL levels (Fig. 4J).

**Fig. 4.**
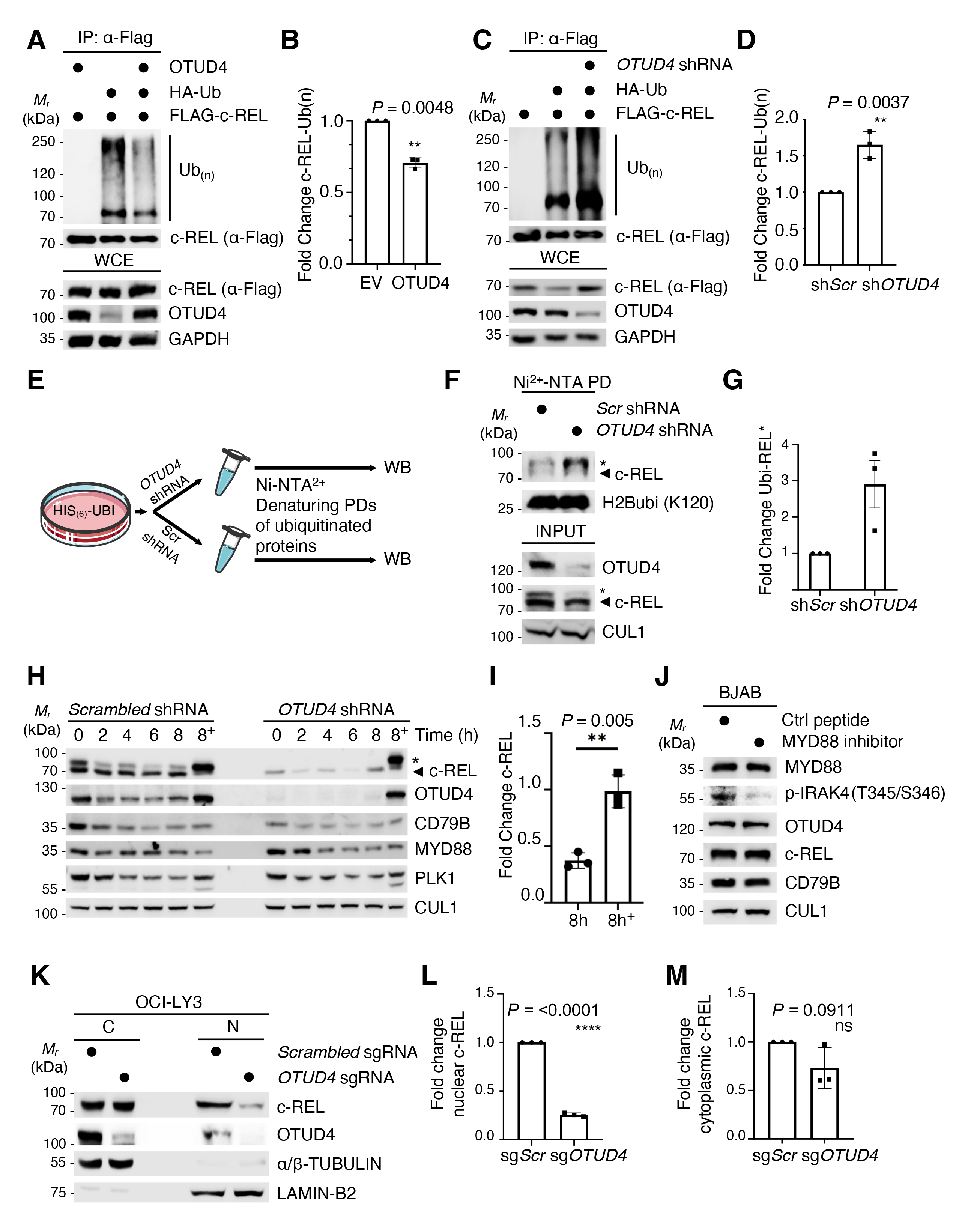
OTUD4 inhibits c-REL degradation and allows c-REL nuclear translocation. **A)** OTUD4 deubiquitination of c-REL. FLAG-c-REL was co-overexpressed with HA-tagged ubiquitin and immunoprecipitated under denaturing conditions upon OTUD4 overexpression and blotted against (*α*- Flag) or a ubiquitin antibody as shown. **B)** Quantification of the ubiquitination signal of immunoprecipitated c-REL in A. The error bar represents the mean ± S.D, (n = 3). **C)** Ubiquitination of FLAG-c-REL upon *OTUD4* downregulation. c-REL was co-overexpressed with HA-ubiquitin in HEK293T cells as indicated. IPs were performed under denaturing conditions upon control or *OTUD4* downregulation. Samples of immunoprecipitated c-REL were blotted as shown. **D)** Quantification of the ubiquitination signal of immunoprecipitated c-REL in 3C. The error bar represents the mean ± S.D, (n = 3). **E)** Scheme of denaturing pull-downs (PDs) of ubiquitinated proteins performed from BJAB cells stably expressing HIS_(6x)_-ubiquitin. DLBCL BJAB cells stably expressing HIS_(6x)_-tagged-ubiquitin were treated with scrambled shRNA or *OTUD4* shRNA and pull-downs of ubiquitin were performed from cell lysates using Ni-NTA^2+^ beads under denaturing conditions. The ubiquitinated proteins on the beads were detected by Western blot (WB) analysis. **F)** Ubiquitination of c-REL in presence and absence of *OTUD4*. Pull-downs of c-REL from shRNA scrambled or shRNA *OTUD4*-silenced DLBCL BJAB cells expressing His-tagged ubiquitin using Ni-NTA beads. Pulled-down proteins were blotted against ubiquitinated-histone 2B (ubiquitin-K120-H2B) as a positive control, and against c-REL. The asterisk (*) depicts the presence of a band of c-REL with lower mobility than expected (arrow). **G)** Quantification of the lower migrating band of c-REL (*) bound to nickel beads (Ni2^+^-NTA PD) in F. The error bar represents the mean ± S.D, (n = 3). **H)** Time-course stability assay of c-REL performed in the BJAB cells used in E-G. Samples of scrambled shRNA or *OTUD4* shRNA treated cells were collected at the indicated time points under cycloheximide (CHX, 100 µg/ml) treatment and under 4h of proteasomal inhibition (MG132, 10 mM) when indicated (^+^). The asterisk (*) depicts a higher MW band of c-REL (arrow). Cell extracts were blotted against shown antibodies using PLK1 as a positive control. **I)** Quantification of fold change of the c-REL signal in H under scrambled shRNA after 8h or CHX treatment in absence and presence (+) of MG132. Significance is included (n=3). **J)** BJAB cells were treated with a control peptide or with MYD88 inhibitor (NBP2-29328, Novus Biologicals) for 48h. WBs using the indicated antibodies revealed the effect of MYD88 inhibition on the depicted proteins. **K)** OTUD4-dependent nuclear localization of c-REL in DLBCL OCI-LY3 cells. Control scrambled sgRNA or *OTUD4* sgRNA depleted OCI-LY3 cells were fractionated into cytoplasmic (C) or nuclear (N) extracts and blotted against the depicted antibodies. **L)** Quantification of the nuclear c-REL levels in K. The error bar represents the mean ± S.D, (n = 3). **M)** Quantification of the cytoplasmic c-REL levels in K. The error bar represents the mean ± S.D, (n = 3).

In line with the finding that the absence of the E3 ubiquitin ligase Peli1 leads to the accumulation of c-Rel in the nucleus of T cells^37^, we hypothesized that deubiquitinase inhibition might have the opposite effect and trigger the absence of c-REL from the B-cell nucleus. We tested the consequences of *OTUD4* silencing in the nuclear localization of c-REL in ABC-type DLBCL cells that show comparable amounts of cytoplasmic and nuclear c-REL. As expected, we found a significant decrease of c-REL in the nuclear fraction of ABC-type OCY-LY3 cells upon inhibition of *OTUD4*, while the cytoplasmic fraction was less affected (Figs. 4K-M). Together, these results identify OTUD4 as a deubiquitinase that counteracts proteasomal degradation of c-REL and allows for its nuclear translocation.

### c-REL-OTUD4 axis is required for the stabilization of BCR and the expansion of DLBCL cells

To confirm the c-REL dependence of CD79B destabilization observed in *OTUD4* downregulated BJAB cells (Fig. 4H), we performed rescue experiments using a doxycycline-inducible over-expression system in BJAB cells. By induced over-expression of c-REL on previously downregulated *OTUD4* cells, CD79B and OTUD4 levels were prominently rescued (Fig. 5A). Conversely, induced over-expression of OTUD4 partially counteracted CD79B and c-REL downregulation upon *c-REL*-targeted shRNA treatment (Fig. 5B) most probably by inhibiting the proteasomal degradation of c-REL amounts still present in the cells. The higher efficacy of over-expressed c-REL in restoring CD79B levels compared to OTUD4 over-expression suggests that within the c-REL-OTUD4 axis c-REL-induced transcriptional signatures have an important contribution to the BCR stabilization. We therefore analyzed the RNA-seq data from our previous *c-REL* downregulation experiments (Fig. 1B) in terms of enrichment analysis to find out which biological functions or pathways are significantly associated with differential expressed genes in *c-REL* downregulated BJAB cells *vs* control cells. Among the 12.251 genes co-expressed in all tested conditions (Supplementary Fig. 5A; Supplementary Table 5) we found a strong correlation of c-REL to genes involved in the regulation of *adaptive immune response* by Gene Ontology (GO) annotation (GO: 0002819) in good accordance with previous bibliography^21, 22, 24^ and interestingly, in *Negative regulation of adaptive immune response based on somatic recombination of immune receptors built from immunoglobulin superfamily domains* (GO: 0002823) (Supplementary Fig. 5; Supplementary Table 5).

**Fig. 5.**
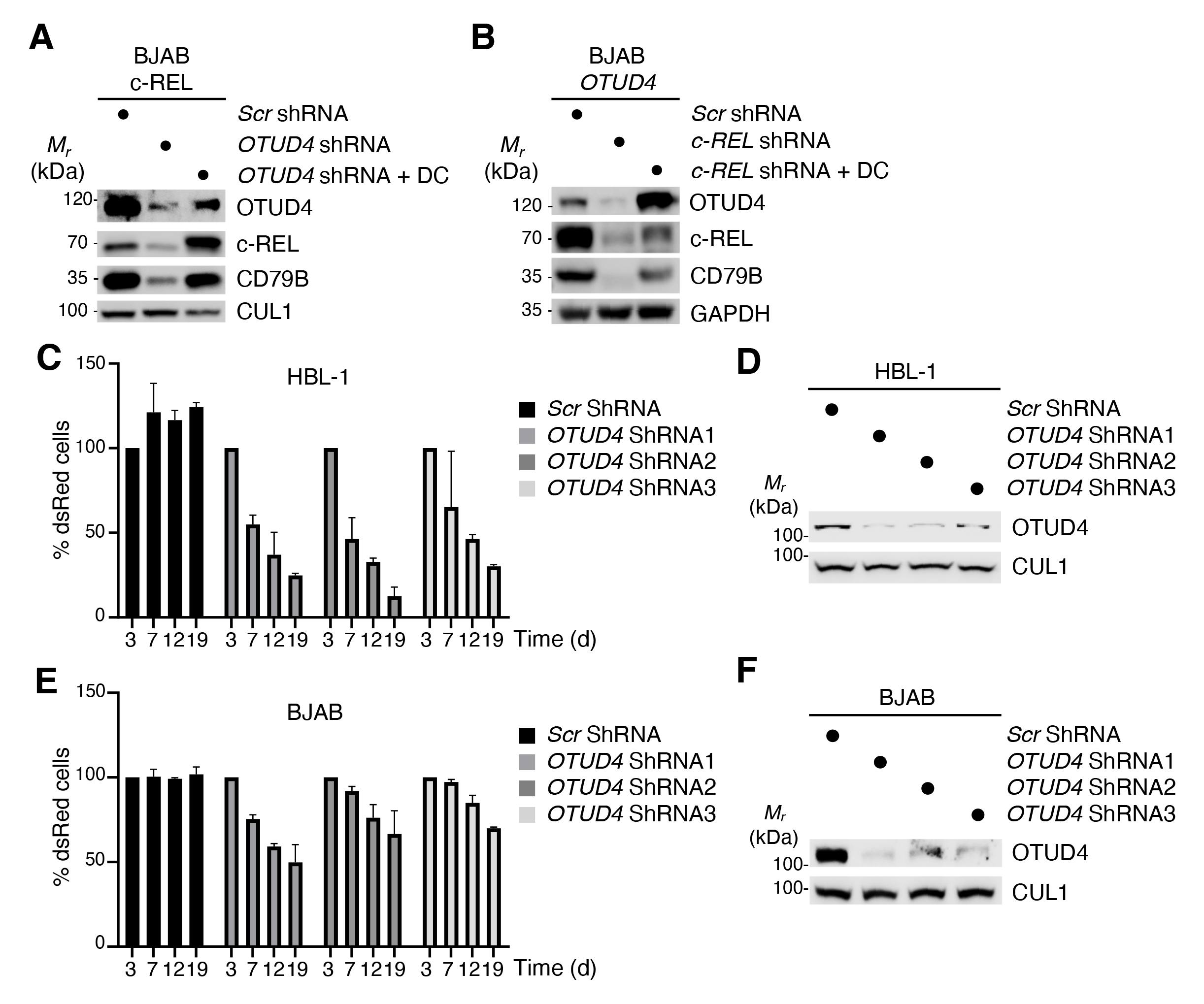
OTUD4 is a vulnerability to destabilize NF-kB and BCR in B-cell lymphoma. **A)** c-REL rescue experiment on *OTUD4* downregulated BJAB cells. BJAB cells stably infected with pTRIPZ-cREL vector under doxycycline induction of expression were treated with scrambled shRNA or *OTUD4* shRNA for 48h. Later, doxycycline (DC) was added as indicated for 24h (1ng/ml). WBs of OTUD4, c-REL, CD79B and CUL1 are shown in the figure. **B)** OTUD4 rescue experiment on c-REL downregulated BJAB cells. A similar experiment to A was performed in BJAB cells stably infected with pTRIPZ-OTUD4 vector under doxycycline induction. Cells were treated with scrambled shRNA or *c-REL* shRNAs for 48h. Later, expression of OTUD4 was induced with doxycycline (DC) as indicated for 24h (1ng/ml). **C)** Cell growth competitive assays of control or *OTUD4*-downregulated HBL-1 cells measured by Fluorescence Assorted Cell Sorting (FACs). Proportion of DLBCL ABC-type HBL-1 cells infected with a control shRNA or *OTUD4* shRNAs (co-expressing dsRed) in competitive co-cultures with unmodified HBL-1 cells. The error bar represents the mean ± S.D, (n = 3). **D)** Western blot showing efficient knock-down of *OTUD4* in the HBL-1 cells used in 5A. **E, F)** Similar experiment as in 5A and 5B performed in DLBCL GC-type BJAB cells.

Downregulation of components of the BCR and NF-κB have been extensively described to impact the proliferation and survival of DLBCL cells^7, 23, 54^. We therefore aimed to decipher the contribution of *OTUD4* to B-cell lymphoma cell expansion. Contrary to GCB-type DLBCL cells, viability of ABC-type DLBCL cells depends on chronic NF-κB active signaling^6^. To determine the biological consequences of *OTUD4* downregulation we silenced *OTUD4* using three different shRNAs and analyzed by flow cytometry the competitive expansion of ABC-type DLBCL HBL-1 cells or GCB-type BJAB cells. Consistent with the role of tonic BCR-NF-κB signals in ABC-type *vs* GCB-type cell, *OTUD4* silencing severely compromised the expansion of ABC-type-HBL-1 cells and, albeit to a lesser extent, of GCB-type BJAB cells (Figs. 5C-F). Together, these data reveal the c-REL-OTUD4 axis for stabilizing the BCR and distinguish OTUD4 deubiquitinase as a dependency of B-cell lymphoma cell expansion.

## Discussion

We discover the c-REL-OTUD4 axis to stabilize BCR subunits in B-cell lymphoma. Thus, we reveal a positive feedback of NF-κB to BCR and fill a gap in the knowledge of the interplay and signaling of NF-κB-BCR in both directions. With the identification of OTUD4 as a vulnerability in B-cell lymphoma that regulates proto-oncogenic c-REL nuclear translocation and the levels of BCR subunits, we address an urgent need to find strategies in the specific regulation of the hitherto undruggable NF-κB transcription factors that frequently lead to relapse in immuno-oncology. Ubiquitination is a reversible process and DUBs are mainly cysteine proteases whose inhibition can induce the degradation of specific proteins, as transcription factors, that are otherwise often undruggable. The importance of our findings is enhanced given the dependency on OTUD4 for B-cell lymphoma cells expansion and the OTUD4 correlation with c-REL in patient samples.

c-REL is a proto-oncogenic protein required for the maintenance of immune checkpoints in cancer^21, 22^ whose levels and nuclear accumulation compared to other transcription factors are primarily mobilized and critical in DLBCL, an entity in which aberrant activation of BCR and *REL* amplification and gain of function occur frequently^13, 29, 52^. The disclosed role of c-REL for the maintenance of the subunit levels of the BCR might be relevant to understand the shift to c-REL-dominated NF-κB heterodimers in mature B cells, and the selective dependency of c-REL in the last stages of B-cell development and antibody responses^20, 52, 55–57^. Our findings call for a systematic investigation of the consequences of c-REL inhibition at different stages of B-cell maturation in relation to BCR stability. Even though the discovery and naming of *Nuclear factor kappa-light-chain-enhancer of activated B cells* (NF-κB) was due to the observation of nuclear proteins that bind the *k* light chain of antibodies^58^, the role of NF-κB transcription factors in the stabilization of the BCR has remained undeciphered. Remarkably, we do not observe direct c-REL transcriptional control of the protein blocks that configure the BCR, rather indirectly through as yet not completely understood multi-feedback loops that regulate the levels of BCR subunits. We reveal a positive feedback loop between the c-REL transcriptional control of deubiquitinase OTUD4, and the OTUD4-mediated stabilization of c-REL protein levels. OTUD4 is a member of the ovarian tumor (OTU) family of deubiquitinases present in human cells that have revealed crucial to regulate cell-signaling cascades^59^. To the best of our knowledge, OTUD4 is the first DUB described to inhibit the ubiquitination and proteasomal degradation of NF-κB c-REL. Previously, OTUD4 has been identified as a dual K63- K48- ubiquitin chain protease to inhibit Rel-A translocation in Mouse Embryonic Fibroblasts^48^, the regulation of alkylation damage response^60^, RNA-binding^53^ and antiviral response^61^. In addition, mutations in *OTUD4* have been described to cause ataxia with hypogonadotropic hypogonadism in humans^62^ and to promote the cell surface levels of TGFβ receptor I^63^. Because c-REL and OTUD4 regulate each other, it will be challenging to dissect and quantify their specific contribution to the stabilization of BCR. Although we have found c-REL to impact at the transcriptional level cellular pathways on somatic recombination of immune receptors, the identification of the precise players modulated by the c-REL-OTUD4 axis to stabilize BCR subunits requires further investigation. In a similar way, additional experiments are required to understand the contrary roles of OTUD4 to inhibit the translocation of Rel-A under NF-κB activation in MEFs^64^, and to allow for c-REL nuclear translocation under non-stimulated conditions in DLBCL (this study). Of notice, OTUD4 has been found as a downstream phosphorylation target of BCR signalosomes^65^. A detailed characterization of the PTMs that regulate OTUD4 activity under different stimuli and their impact on NF-κB modulation will be of most interest.

The emerging picture of NF-κB-BCR interrelation based on the results reported by others^3, 26–28^ and our study prompt us to propose a model by which in addition to IκBs-mediated cytoplasmic retention of NF-κB subunits, OTUD4 deubiquitination and stabilization of c-REL exerts further control to generate NF-κB activity that is auto-regulated by the transcriptional control of *OTUD4* expression in a feedback loop mechanism, that finally regulates the levels of BCR subunits in a still undeciphered manner (Fig. 6).

**Fig. 6.**
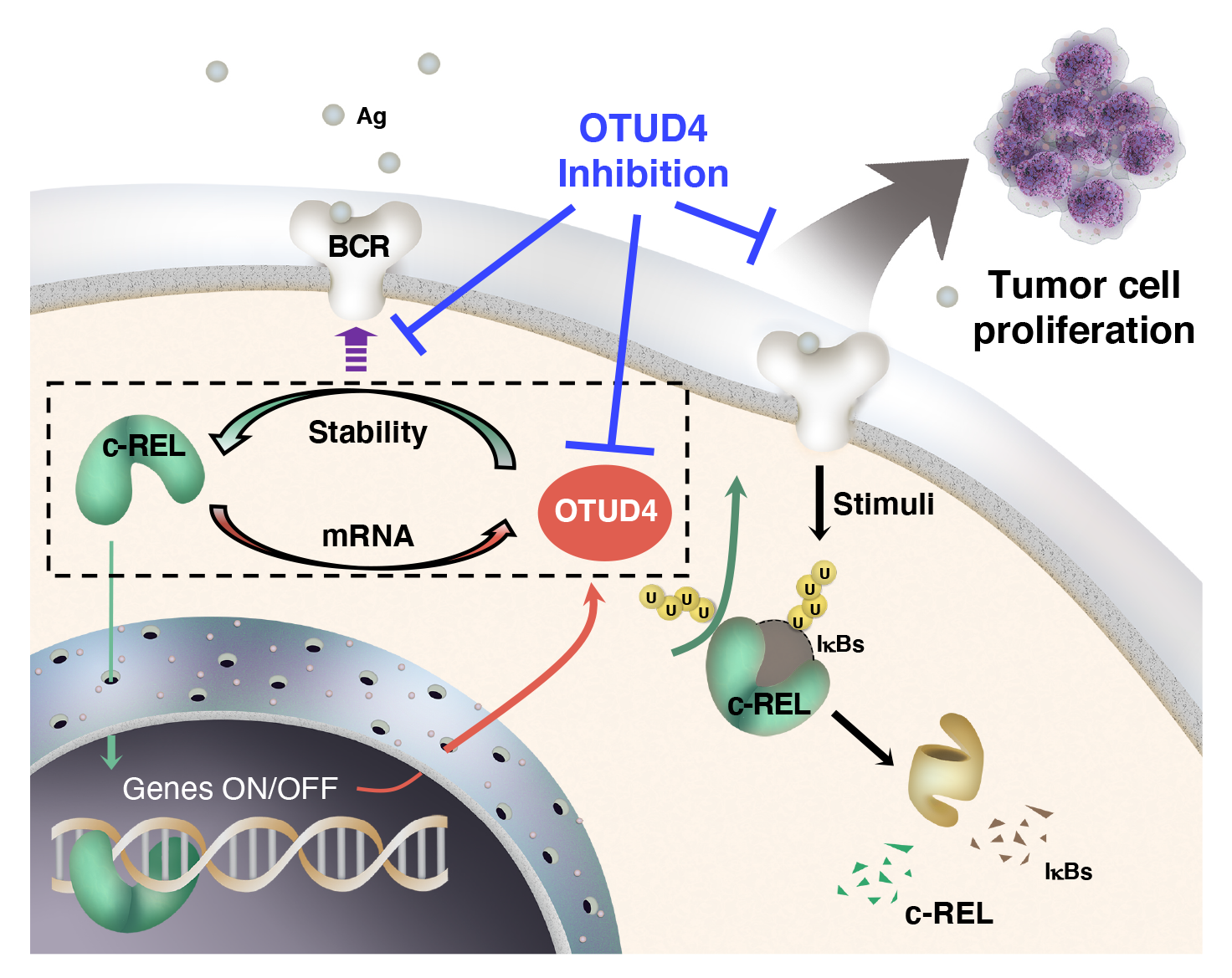
NF-κB c-REL-OTUD4 axis in B-cell lymphoma. Upon stimuli, in addition to protein turnover of the NF-κB inhibitory proteins (IκBs) to allow nuclear translocation of NF-κB subunits, we propose a further control of free NF-κB abundance mediated by direct de-ubiquitination and recovery of c-REL transcription factor otherwise targeted for proteasomal degradation. The additional control mediated by OTUD4 deubiquitination of c-REL, enables translocation of the proto-oncogenic NF-κB subunit to the nucleus. Subsequent induction of a malignant transcriptional program includes *OTUD4* mRNA regulation and BCR stabilization in a multi-loop feedback mechanism of c-REL to BCR. Thefore, the axis formed by c-REL-OTUD4 is required for the stabilization of BCR and cancer cell growth. In our model, OTUD4 inhibition might result in synergistic blockage of NF-κB c-REL and BCR signaling pathways.

In summary, our findings provide new rationale to synergistically inhibit the NF-κB c-REL transcription factor and BCR signaling in c-REL-mediated auto-immune diseases and in B-cell lymphoma. Contrary to the current bulk of clinical drugs such as bortezomib and immunomodulatory drugs (IMiDs) that among other broad consequences also target pathological NF-κB signaling^66, 67^, OTUD4 inhibition could provide a selective mode of action on enhanced c-REL with a narrower biological effect and undesired NF-κB-mediated drug resistance in BCR-targeted therapies.

## Methods

### Cell lines and drug treatments

HEK 293T (ATCC: CRL-3216) were grown in Dulbecco’s modified Eagle’s medium (DMEM; Gibco) supplemented with 10% fetal calf serum (FCS) and 1% penicillin-streptomycin. The human DLBCL cell lines were cultured in RPMI-1640 (Gibco) with 15% heat-inactivated fetal bovine serum (FBS) and 1% penicillin-streptomycin. All cells tested negative for mycoplasma by a PCR detection method. DLBCL cell lines BJAB, HBL-1, OCI-LY3 and TMD8 were further authenticated. MG132 (TOCRIS, 1748), cycloheximide (Sigma-Aldrich, C7698), doxycycline (Sigma-Aldrich, D5207) and MYD88 inhibitor (Novus, NBP2-29328) were added according to the manufacturers’ instructions.

### Plasmids, shRNAs and sgRNAs

Human c-REL (isoform NP_002899.1) template was described previously^52^. c-REL was cloned into pcDNA3.1 (Life technologies) and pTRIPZ (Horizon) using the following oligos: forward (NheI) 5´- GCG GCT AGC GCC TCC GGT GCG TAT AAC CCG - 3´ and reverse (XhoI) 5´- GCC GAG CTC TTA TAC TTG AAA AAA TTC −3´. Human *OTUD4* (isoform 4 NP_001352986.1) was purchased from GenScript. Human *OTUD4* constructs were cloned without tag or with FLAG-tag into the expression plasmid pcDNA3.1 (Life technologies) using the oligos forward (BamHI) 5´- CGG GGA TCC ATG GAG GCT GCC GTC GGC GTC CC - 3 and reverse (XhoI) 5´- CGG CTC GAG TCA AGT GTG CTG TCC CCT ATG −3´. pRK5-HA-Ubiquitin-WT was obtained from Addgene (Addgene #17608). pTRIPZ shRNA vectors were purchased from Horizon. The HIS_(6x)_-ubiquitin in pLenti for stable expression of HIS_(6x)_- tagged-ubiquitin was described previously^68^. For shRNA mediated silencing of c-REL, specific shRNAs were cloned into the pLKO.1 TRC plasmid (Addgene #10878), where the puromycin resistance cassette was replaced by the cDNA coding for DsRed-Express2. The shRNA target sequences for c-REL were: (#1) 5´-CCG GAC AAA GAA TGA CCC ATA TAA ACT CGA GTT TAT ATG GGT CAT TCT TTG TTT TTT G-3´, (#2) 5´-CCG GTG TTG TCT CGA ACC CAA TTT ACT CGA GTA AAT TGG GTT CGA GAC AAC ATT TTT G-3´and (#3) 5´-CCG GCA GAG GAG GAG ATG AAA TAT TCT CGA GAA TAT TTC ATC TCC TCC TCT GTT TTT G-3´. For shRNA mediated silencing of *OTUD4*, specific shRNAs were cloned into the pLKO.1 TRC plasmid (Addgene #10878), where the puromycin resistance cassette was replaced by the cDNA coding for DsRed-Express2. Sh_scramble (Addgene #10879) was used as control. The shRNA target sequences for *OTUD4* were: (#1) 5´- TGCTGTATGAGAAGGTATTTA −3´, (#2) 5´- GAATCCAAGCAAGCCAATAAA −3´ and 5´- CTGTATCCCAAGCTCATTTAA −3´ for *CTRL* / *sh*_*scramble were*: 5´- CCTAAGGTTAAGTCGCCCTCG −3’. Depletion of *OTUD4* by CRISPR Cas9 was performed by cloning specific sgRNA into lentiCRISPRv2 (Addgene #98290) as described in Wang *et al.*, 2015. sgRNA sequences used were 5´- CGCTTCCGCGGCCCGTTCAA −3´ for scramble sgRNA and 5-ACAACAGATGTGGATTACAG −3´ for *OTUD4* sgRNA.

### Antibodies, cell lysis, immunoprecipitations and immunoblotting

*Antibodies*. The following primary antibodies were used for Western blotting: α/β-Tubulin (1:1,000, rabbit, Cell Signaling Technology #2148), c-REL (1:1,000, goat, R&D Systems #AF2699), c-REL (1:1,000, rabbit, Cell Signaling Technology #4727), CUL1 (1:500, mouse, Sigma-Aldrich #32-2400), CD79B (1:1000, rabbit, Cell Signaling Technology #96024), FLAG (1:1,000, rabbit, Sigma #F7425), GAPDH (1:1,000, mouse, Santa Cruz #sc-47724), GST (1:1,000, mouse, Santa Cruz #sc-138), HA (1:1,000, rabbit, Cell Signaling Technology #3724), IkBa (1:1,000, mouse, Cell Signaling Technology #4814), Lamin B2 (1:1,000, rabbit, Cell Signaling Technology #13823), MyD88 (1:1,000, mouse, Cell Signaling Technology #50010), OTUD4 (1:1,000, rabbit, Sigma-Aldrich #HPA036623), OTUD6B (1:1,000, rabbit, Abcam #ab127714), PLK1 (1:500, mouse, Thermo Fisher # 33-1700), Ubiquitin FK2 (1:1,000, mouse, Enzo #BML-PW8810-0100). Secondary antibodies (anti-rabbit IgG, anti-mouse IgG or protein-A) coupled with horseradish peroxidase were from GE Healthcare, an anti-goat IgG secondary antibody was from Santa Cruz.

### Cell lysis, pull-downs, immunoprecipitations and immunoblotting

Cell samples were lysed in standard 150mM NaCl lysis buffer (NaCl 150mM, Tris-HCl 50mM pH 7.5, MgCl_2_ 5mM, EDTA 1mM, NP-40 0.1%, Glycerol 5% and protease inhibitors) unless otherwise specified. Extract preparation, immunoprecipitation, and immunoblotting have been previously described^69^. The cellular lysates were centrifuged 15 min at 14,000 rpm at 4°C to separate from cellular debris, protein concentration was measured using a Bio-Rad DC protein assay (Lowry assay) and Laemmli buffer was added. The lysates were separated by SDS-PAGE and blotted onto PVDF membranes (Millipore). Equal protein levels were confirmed by Ponceau S staining. After blocking unspecific binding in 5% milk, membranes were incubated with primary antibodies diluted in 5% milk or 5% bovine serum albumin (BSA). After addition of horseradish peroxidase coupled secondary antibodies, Western blot membranes were developed using the enhanced chemo luminescence (ECL) method (SuperSignal West, Thermo Fisher). Visualization of the protein bands was performed using the Fusion FX imaging system (VILBER), and the intensity of the signal was quantified by the accompanying software Evolution Capt V18.02c (VILBER).

For immunoprecipitations (IPs), cell lysates were incubated with beads overnight. For endogenous IPs, WCE were subjected to a pre-clear with Protein A beads. After washing the beads four times in lysis buffer, Laemmli buffer was added and proteins were analyzed by SDS-electrophoresis.

#### Cellular fractionation

To purify target proteins from cytoplasmic and nuclear compartments, the NE-PERTM Nuclear and Cytoplasmic Extraction kit (Thermo Fisher, 78835) was used. Cells were freshly harvested and processed according to the manufacturer’s protocol. Obtained cytoplasmic and nuclear fractions were either directly separated on SDS-PAGE after denaturing by cooking with Laemmli buffer at 95°C or frozen at −20°C till further use.

#### Tandem Ubiquitin Binding Entities (TUBEs) and Ni-NTA^2+^-agarose Pull-downs

Cells were first fractionated using the Nuclear and Cytoplasmic Extraction kit (Thermo Scientific). The resultant cytoplasmic and nuclear lysates were incubated with the corresponding beads for 2 h rotating at 4°C. After four consecutive washes with 150Mm NaCl lysis buffer by centrifugation at 1,200 rpm for 1 minute, the precipitated beads were boiled for 10 minutes with 5x Laemmli buffer followed by immunoblotting. For the Ni-NTA^2+^-agarose (Qiagen #1018244) pull-downs, denaturing conditions were employed following manufacturés protocol.

### Transient transfections and lentivirus-mediated DNA transfer

HEK 293T cells were transfected using the calcium phosphate method: A 1 mL transfection mixture was prepared by dissolving 10 µg of DNA in 450 µL dH2O with the addition of 50 µL CaCl_2_ to a final concentration of 250mM. Then, 500 µL 2x BES solution (Sigma, 14280) was added dropwise to the tube walls with rotation. The mixture was left at room temperature for 20 minutes and later dropped carefully onto the cells while swirling the plate. The medium was replaced with fresh medium after 4 h.

Lentiviral particles were produced in HEK293T cells by transfecting a 10 cm plate with 15 µg packaging plasmid psPAX2(Addgene), 5 µg envelope plasmid pMD2.G (Addgene), and 20 µg transfer plasmid (pTRIPZ, pLKO.1 dsRed plasmid for shRNA and lentiCRISPRv2 for sgRNA) using the aforementioned calcium phosphate method. After 4 h, the medium was replaced with 10 mL of fresh medium, and the transfected cells were incubated for 48 h. The viral supernatant was harvested on the 2^nd^ day by passing the collected supernatant through a 0.45 µm filter and directly used for infection or stored at −80°C.

For infection, DLBCL cell lines or HEK293T cells were plated in 6 well plates and incubated with the virus-containing supernatant for 24 hours. DLBCL cells were spin-infected at 700 g for 30 min at 37°C.

### In vivo deubiquitylation

HEK293T cells were transiently transfected with 1 µg of HA Ubiquitin, 5 µg of FLAG-c-REL (substrate) and 5 µg of OTUD4 (DUB). Twenty-four hours after transfection, the cells were harvested and lysed in 100 µL of 150mM NaCl lysis buffer supplemented with inhibitors and incubated on ice for 30 minutes. After centrifugation at 14,000 rpm speed for 20 minutes, the lysates were transferred into new Eppendorfs, denatured with 10 µL 10% SDS and 1 µL EDTA (0.5 M), and cooked at 95°C for 5 minutes. The samples were then cooled down to room temperature for 5 minutes before quenching with 900 µL of 1% Triton X-100. Later, the mixture was cooled on ice for 5 minutes, followed by the addition of 25µL pre-washed FLAG-beads and processed for WB as discussed before.

#### Quantitative proteomics

For total proteome analysis, BJAB cells were lysed in 2% SDS and 50 mM Tris-HCl pH 7.5 and heated to 95°C for 10 min. 1 µl 100% TFA was added to each sample to hydrolyze DNA and the pH was subsequently adjusted to 8.5 with 3 M Tris solution. Detergent was removed from lysates by SP3 cleanup, following the protocol first described by Hughes et al. (PMID: 30464214) Briefly, lysate containing 15 µg of protein was mixed with SP3 beads, and proteins were precipitated onto a 50:50 mixture of Sera-Mag Speed Bead types A and B (Thermo Fisher Scientific) in 70% acetonitrile. Beads were washed three times with 80% ethanol in water and once with acetonitrile. Disulfide bond reduction was performed in 50 µl 50 mM Tris-HCl pH8.5 with 10 mM DTT for 45 min at 30°C, followed by alkylation of cysteines with 55 mM CAA for 30 min at room temperature. Tryptic digest was performed by adding 50 µl of digestion buffer (2 mM CaCl_2_ in 50 mM Tris-HCl pH 8.5) with a 1:25 enzyme-to-protein ratio and digest at 37°C overnight. On the next day, the supernatant of each sample was transferred to a new vial and acidified with 1% formic acid (FA) followed by desalting on self-packed StageTips (three disks, Ø 1.5 mm C18 material, 3 M Empore^TM^). Peptides were eluted with 50% ACN, 0.1 % FA and dried down prior to MS analysis.

#### MS analysis

Peptides were reconstituted in 0.1 % formic acid (FA) and analyzed by nanoLC-MS/MS on a Dionex Ultimate 3000 UHPLC+ system coupled to a Q Exactive HFX mass spectrometer (Thermo Fisher Scientific). After 10 min of washing (0.1 % FA, 5 µL/min) on a trap column (75 um x 2 cm, 5 µm C18 resin; Reprosil PUR AQ, Dr. Maisch), peptides were transferred to an analytical column (75 µm x 45 cm, 3 µm C18 resin; Reprosil Gold, Dr. Maisch) and separated at 300 nL/min using a 50 min linear gradient from 4 % to 32 % LC solvent B (0.1 % FA, 5 % dimethyl sulfoxide (DMSO) in acetonitrile) in LC solvent A (0.1 % FA in 5 % DMSO). The Q Exactive HFX was operated in data dependent and positive ionization mode. Full-scan mass spectra (m/z 360-1300) were acquired in profile mode with 60,000 resolution, an automatic gain control (AGC) target value of 3e6 and 45 msec maximum injection time. For the top 18 precursor ions, high resolution MS2 scans were performed using HCD fragmentation with 26% normalized collision energy, 15,000 resolution, an AGC target value of 1e5, 25 msec maximum injection time and 1.3 *m/z* isolation width in centroid mode. The intensity threshold was set to 8.8e4 with a dynamic exclusion of 25sec.

Peptide and protein identification and quantification for the full proteome samples were performed using MaxQuant 2.0.3.0 by searching the MS2 spectra against the human reference proteome supplemented with common contaminants. Match-between-runs, iBAQ (intensity based absolute protein quantification) computation, and label-free quantification were enabled. All other search parameters were left as default. Hits to the reverse and contaminant databases were removed and LFQ values used for proteome comparison. Protein abundance values were loaded onto Perseus 1.6.14.0 and further processed. Briefly, abundance values were log2 transformed and only proteins with at least 3 valid values in at least one of the groups under analysis were considered for further processing. Then, missing values were replaced with values from the normal distribution, using a width of 0.1. Finally, a t-test was applied to compare scrambled control and targeted-gene shRNA samples and proteins with a p value<0.05 and log2 fold change smaller −0.5 or larger 0.5 were considered as significantly different.

#### MS data availability

The MS proteomics raw data and MaxQuant search results have been deposited to the ProteomeXchange Consortium (http://www.proteomexchange.org/) via the PRIDE partner repository with the data set identifier 10.6019/PXD041576.

### RNA-seq

Total RNA was extracted and purified from transfected BJAB cells using RNeasy Plus Kits (QIAGEN #74034) containing gDNA eliminator columns. With extracted total RNA, RNA-seq library was constructed. Messenger RNA was purified from total RNA using poly-T oligo-attached magnetic beads. After fragmentation, the first strand cDNA was synthesized using random hexamer primers, followed by the second strand cDNA synthesis using either dUTP for directional library or dTTP for non-directional library. For the non-directional library, it was ready after end repair, A-tailing, adapter ligation, size selection, amplification, and purification. For the directional library, it was ready after end repair, A-tailing, adapter ligation, size selection, USER enzyme digestion, amplification, and purification. The library was checked with Qubit and real-time PCR for quantification and bioanalyzer for size distribution detection. Quantified libraries will be pooled and sequenced on Illumina platforms, according to effective library concentration and data amount. Clustering and sequencing. The clustering of the index-coded samples was performed according to the manufacturer’s instructions. After cluster generation, the library preparations were sequenced on an Illumina platform and paired-end reads were generated.

#### Data Analysis

Quality control. Raw data (raw reads) of fastq format were firstly processed through in-house perl scripts. In this step, clean data (clean reads) were obtained by removing reads containing adapter, reads containing ploy-N and low-quality reads from raw data. At the same time, Q20, Q30 and GC content the clean data were calculated. All the downstream analyses were based on the clean data with high quality. Reads mapping to the reference genome. Reference genome and gene model annotation files were downloaded from genome website directly. Index of the reference genome was built using Hisat2 v2.0.5 and paired-end clean 2 reads were aligned to the reference genome using Hisat2 v2.0.5. We selected Hisat2 as the mapping tool for that Hisat2 can generate a database of splice junctions based on the gene model annotation file and thus a better mapping result than other non-splice mapping tools.

Quantification of gene expression level featureCounts v1.5.0-p3 was used to count the reads numbers mapped to each gene. And then FPKM of each gene was calculated based on the length of the gene and reads count mapped to this gene. FPKM, expected number of Fragments Per Kilobase of transcript sequence per Millions base pairs sequenced, considers the effect of sequencing depth and gene length for the reads count at the same time, and is currently the most commonly used method for estimating gene expression levels. Differential expression analysis (For DESeq2 with biological replicates) Differential expression analysis of two conditions/groups (two biological replicates per condition) was performed using the DESeq2 R package (1.20.0). DESeq2 provide statistical routines for determining differential expression in digital gene expression data using a model based on the negative binomial distribution. The resulting P-values were adjusted using the Benjamini and Hochberg’s approach for controlling the false discovery rate. Genes with an adjusted P-value <=0.05 found by DESeq2 were assigned as differentially expressed. (For edgeR without biological replicates) Prior to differential gene expression analysis, for each sequenced library, the read counts were adjusted by edgeR program package through one scaling normalized factor. Differential expression analysis of two conditions was performed using the edgeR R package (3.22.5). The P values were adjusted using the Benjamini & Hochberg method. Corrected P-value of 0.05 and absolute foldchange of 2 were set as the threshold for significantly differential expression. Enrichment analysis of differentially expressed genes Gene Ontology (GO) enrichment analysis of differentially expressed genes was implemented by the clusterProfiler R package, in which gene length bias wascorrected. GO terms with corrected Pvalue less than 0.05 were considered significantly enriched by differential expressed genes. Gene Set Enrichment Analysis Gene Set Enrichment Analysis (GSEA) is a computational approach to determine if a pre-defined Gene Set can show a significant consistent difference between two biological states. The genes were ranked according to the degree of differential expression in the two samples, and then the predefined Gene Set were tested to see if they were enriched at the top or bottom of the list. Gene set enrichment analysis can include subtle expression changes. We use the local version of the GSEA analysis tool http://www.broadinstitute.org/gsea/index.jsp, GO、KEGG、Reactome、DO and DisGeNET data sets were used for GSEA independently.

### Chromatin immunoprecipitation Sequencing (ChIP-seq)

*ChIP*. Cells were seeded onto the cell culture dishes on the day before the experiment. Next day, cell culture media were replaced by the media without FBS and cells were fixed using 0,8% formaldehyde for 10 min at RT. Quenching was performed adding 125mM glycine by shaking at RT for 5 min. Cells were washed with PBS, collected, snap frozen and kept at −80°C. Cell pellets were resuspended in Farham buffer (5 mM PIPES, pH 8; 85 mM KCl; 0.5% IGEPAL, protease inhibitors) and transferred to the AFA tubes (12×12, Covaris, 520081). Tubes were sonicated at peak power 75 W, duty factor 2% and 200 Cycles/burst at 4°C, for 5 min using Covaris Ultrasonicator E220. Samples were transferred to the new tubes, centrifuged at 1000 g, 4°C for 5 min and supernatants were removed. Pelleted nuclei were washed with Farham buffer, then resuspended with 1 mL chromatin shearing buffer (10 mM Tris-HCl, pH 8; 0.1% SDS; 1 mM EDTA) and transferred to the new AFA tubes. Tubes were sonicated at peak power 140 W, duty factor 5% and 200 Cycles/burst at 4°C, for 15 min using Covaris Ultrasonicator E220. Sheared chromatin samples were transferred to new tubes for centrifugation at maximum speed, 4°C for 10 min. Supernatants were kept at −80°C for further steps. Quality of sheared chromatin was analyzed using TapeStation (Agilent Technologies). 50µL of input from each sample were de-crosslinked using 200 mM NaCl at 65°C for overnight, treated with RNaseA at 37°C for 30 min and Proteinase K at 65°C for 1 h, purified with ChIP DNA Clean & Concentrator™ (Zymo Research, D5205). ChIP was performed using 30 µg chromatin per input sample in ChIP dilution buffer (23 mM Tris-HCl, pH 8; 200 mM NaCl, 2.3 mM EDTA, 1.3% Triton X, protease inhibitors) and c-REL antibody (Cell signalling, 4727, 1:100). Samples were incubated at 4°C overnight and the next day, precipitation was performed using BSA (5mg/mL) pre-blocked protein A and G magnetic beads (Magna ChIP™, 16-663) at 4°C for 2 h. Beads were washed with the following ice-cold buffers ((Low salt buffer (0.1% SDS, 1% Triton X-100, 2 mM EDTA, 20 mM Tris-HCL pH8, 150 mM NaCl), high salt buffer (0.1% SDS, 1% Triton X-100, 2 mM EDTA, 20 mM Tris-HCL pH8, 500 mM NaCl), LiCl buffer (0.25 M LiCl, 1% IGEPAL-CA630, 1% Sodium Deoxychelate, 1 mM EDTA, 10 mM Tris-HCL pH8) and TE buffer (10 mM Tris-HCL pH8, 1 mM EDTA)) twice at 4°C for 10 min each. ChIPed samples were eluted using 150 µL of elution buffer (1% SDS, 50 mM NaHCO_3_) by rotation at RT for 30 min. This step was repeated once. ChIPed samples were de-crosslinked using 200 mM NaCl at 65°C for overnight, treated with RNaseA at 37°C for 30 min and Proteinase K at 65°C for 1 h, purified with ChIP DNA Clean & Concentrator™ (Zymo Research, D5205).

#### ChIPseq library prep and sequencing

ChIPseq libraries were prepared from 1ng ChIPed material (or 3ng for input) of each sample using Ovation® Ultralow V2 DNA-Seq Library Preparation Kit (0344NB, Tecan), according to standard manufacturer’s protocol. Starting materials and the final library quality controls were performed using Qubit™ Flex Fluorometer (Q33327, Thermo Fisher Scientific) and 4200 TapeStation System (G2991BA, Agilent) before and after library preparation. Paired-end sequencing was performed on Illumina NovaSeq 6000 (60bp - 60bp reads). Total samples were multiplexed and sequenced in one lane of SP 100 cycles kit to reduce a sequencing batch effect. BCL raw data were converted to FASTQ data and demultiplexed by bcl2fastq Conversion Software (Illumina). Fastq files were mapped against the humen genome (build GRCh38.p13) using the STAR aligner (version 2.7.10b) using the following parameters: outFilterMultimapNmax 1; alignIntronMax 1; outFilterMismatchNmax 1; alignEndsType EndToEnd to ensure unique mapping. For visualization purposes mapped files were then converted to the bigwig format for visualization using the the bam2wig script from the RSeQC (verson 5.0.1) and the wig2bigwig script from the UCSC repository. Peak calling was done using the MACS3 tool (version 3.0.0b1) with a quality threshold of q<=0.01. Data has been deposited into GEO database with the GSE230160 repository number.

### Flow cytometry

DLBCL cells infected with the dsRed-expressing pLKO.1 plasmid were harvested at the indicated time points, washed twice in PBS and the intensity of the dsRed expression was measured at the PE channel CytoFLEX LX (Beckman Coulter) and the data was analyzed using the FlowJo software (Tree Star Inc, Stanford).

### Immunohistochemistry on Tissue Microarray (TMA)

A TMA from DLBCL patients described previously^70^ was immunostained against validated c-REL (Cell Signaling #4727) and OTUD4 (Sigma #HPA036623) antibodies (1:100). A certified pathologist assessed the expression score of c-REL and OTUD4 in tumour cells according to the following classification; 0 (no expression), 1 (<10%), 2 (10-50%), 3 (51-80%), and 4 (>80%). Scores 0-2 were considered as low and scores 3-4 as high. Informed consent was obtained from patients for the scientific use of biopsy material. The ethics committees of the Technische Universität München approved data analysis (ethics approval 498/17s).

### Quantification and statistical analysis

Statistical analyses of the results were performed by paired and unpaired t-tests, according to assumptions of the test, using GraphPad Prism software as indicated in the figure legends. Unless specified, the error bars shown in the figures and Supplementary Figures represent the mean ± S.D. The *P* values are presented in the figure legends where a statistically significant difference was found: *, *P* < 0.05; **, *P* < 0.01; ***, *P* < 0.001; ****, *P* < 0.0001. n indicates the number of independent biological replicates of an experiment.

## Acknowledgements

We are greatly thankful to Prof. Dr. Marc Schmidt-Supprian and Prof. Dr. Florian Bassermann for equipment and encouraging comments. We thank Dr. Gonca Çetin for preparing the ChIP samples. We thank Jana Zecha, Bernhard Kuster, Maike Kober and Daniel Krappmann for initial help and constructive remarks. We thank Reinhard Fässler and Mercedes Costell for continuous support. This work was supported by Deutsche Forschungsgemeinschaft (DFG; German Research Foundation) (project ID 360372040-SFB 1335 to V.F. S. and K. S.). We thank the *NGS Core Facility at the Max Planck Institute of Biochemistry*, the *Bavarian Center for Biomolecular Mass Spectrometry (BayBioMS) of the TUM*, and the *Tissue bank of Klinikum rechts der Isar and TUM and the Comparative Experimental Pathology* for excellent technical support.

## Author contributions

V. F. S. designed the research; E. K. performed most of the experiments with help from A. J. K.; K. S. performed the IHC analysis of patient samples; J. M. performed the total proteome and analysis; A. Y. performed ChIP-seq and mRNA-seq analysis. M. A. and F. E. performed the multiomic analysis. Results were analyzed by E. K, K. S., J. M., Y. A., M. A., F. E. and V. F. S.; V. F. S. coordinated this work; V. F. S. wrote the manuscript. All authors discussed the results and commented on the manuscript.

---

**Supplementary Fig. 1.**
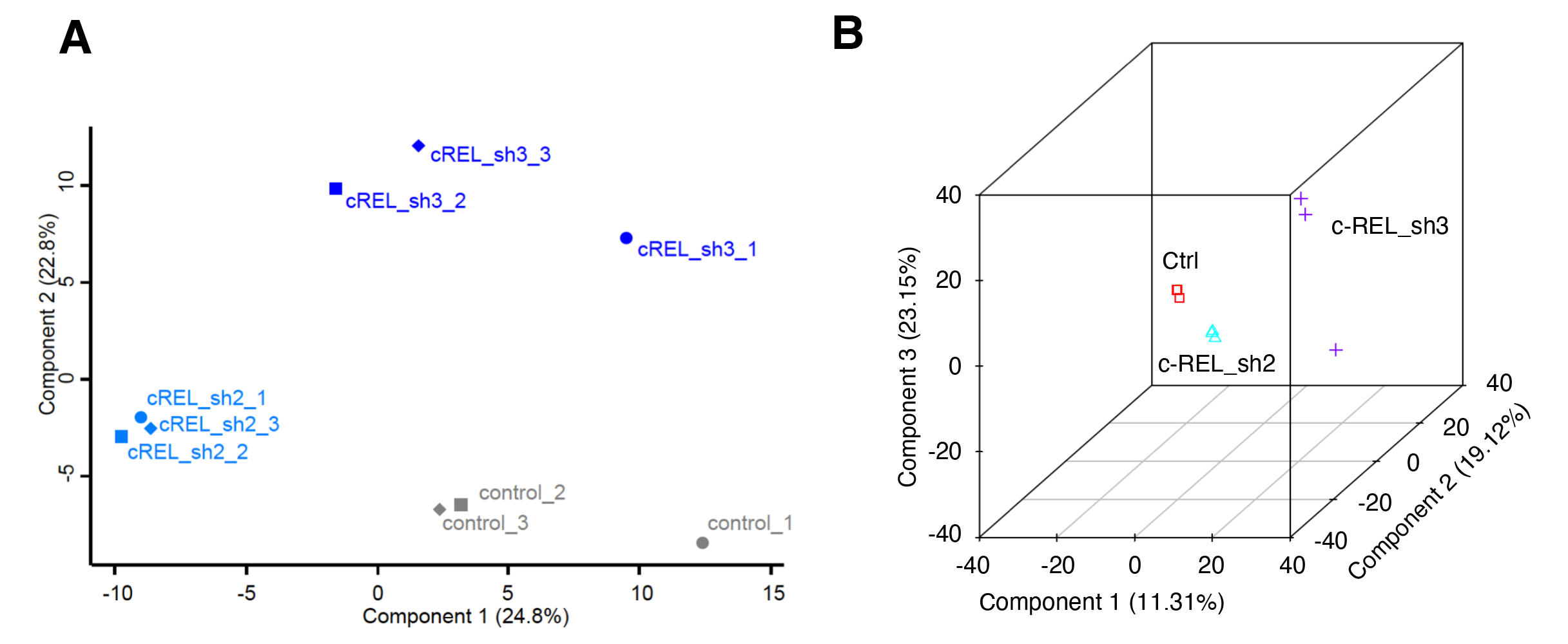
Principal component analysis (PCA) plots. **A)** 2D PCA plot corresponding to MS-data of BJAB cells. Triplicates of control (grey) and two shRNAs targeted against *c-REL* (blue) are depicted in the figure. **B)** 3D PCA plot corresponding to mRNA-Seq analysis of BJAB cells. Triplicates of control (red squares) and two different shRNAs to downregulate c-REL (blue triangles and purple crosses) are depicted in the figure. PCA analysis of gene expression value (FPKM) was performed of all samples and shown in the figure.

**Supplementary Fig. 2.**
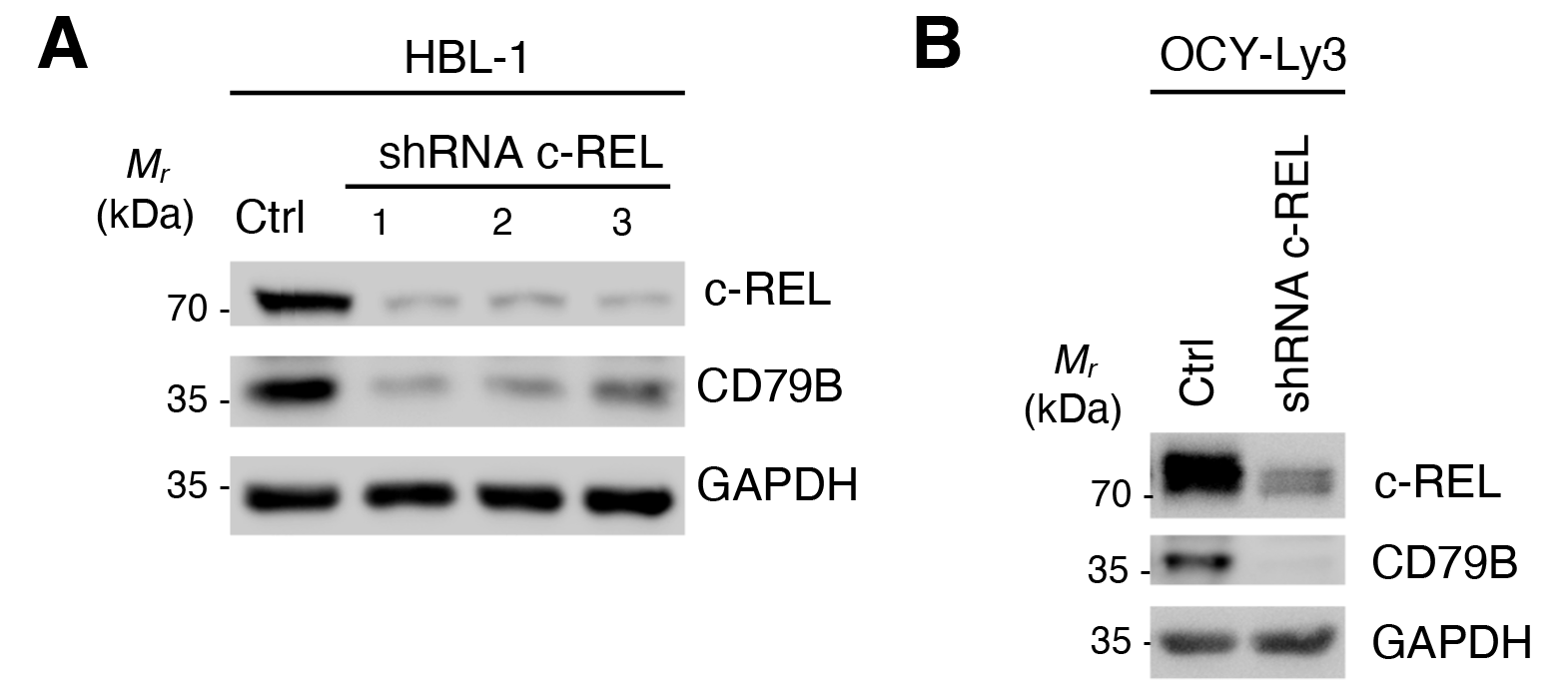
*c-REL* downregulation destabilizes CD79B in DLBCL cells. **A)** Western blot analysis of DLBCL HBL-1 cells upon *c-REL* downregulation using three different shRNAs against *c-REL.* c-REL, CD79B, or GAPDH proteins are blotted against the indicated antibodies. Same experiment as in A was performed in DLBCL OCY-Ly3 cells using shRNA3.

**Supplementary Fig. 3.**
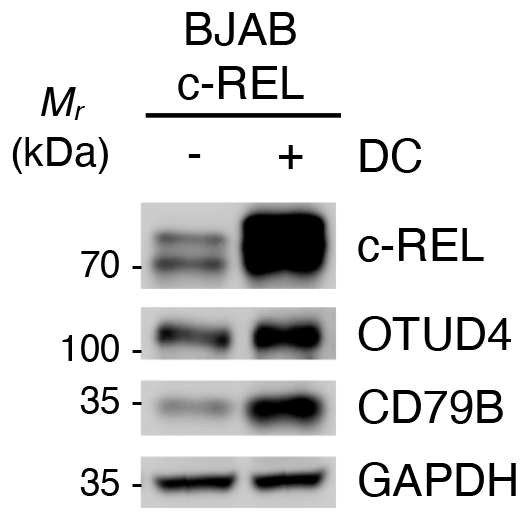
c-REL over-expression regulates OTUD4 expression in BJAB cells. Western blot analysis of BJAB cells upon c-REL over-expression using an inducible expression system in absence (-) or presence (+) of doxycycline (DC) inducer for 24h (1ng/ml). c-REL, OTUD4, CD79B, or GAPDH proteins are blotted against the indicated antibodies.

**Supplementary Fig. 4.**
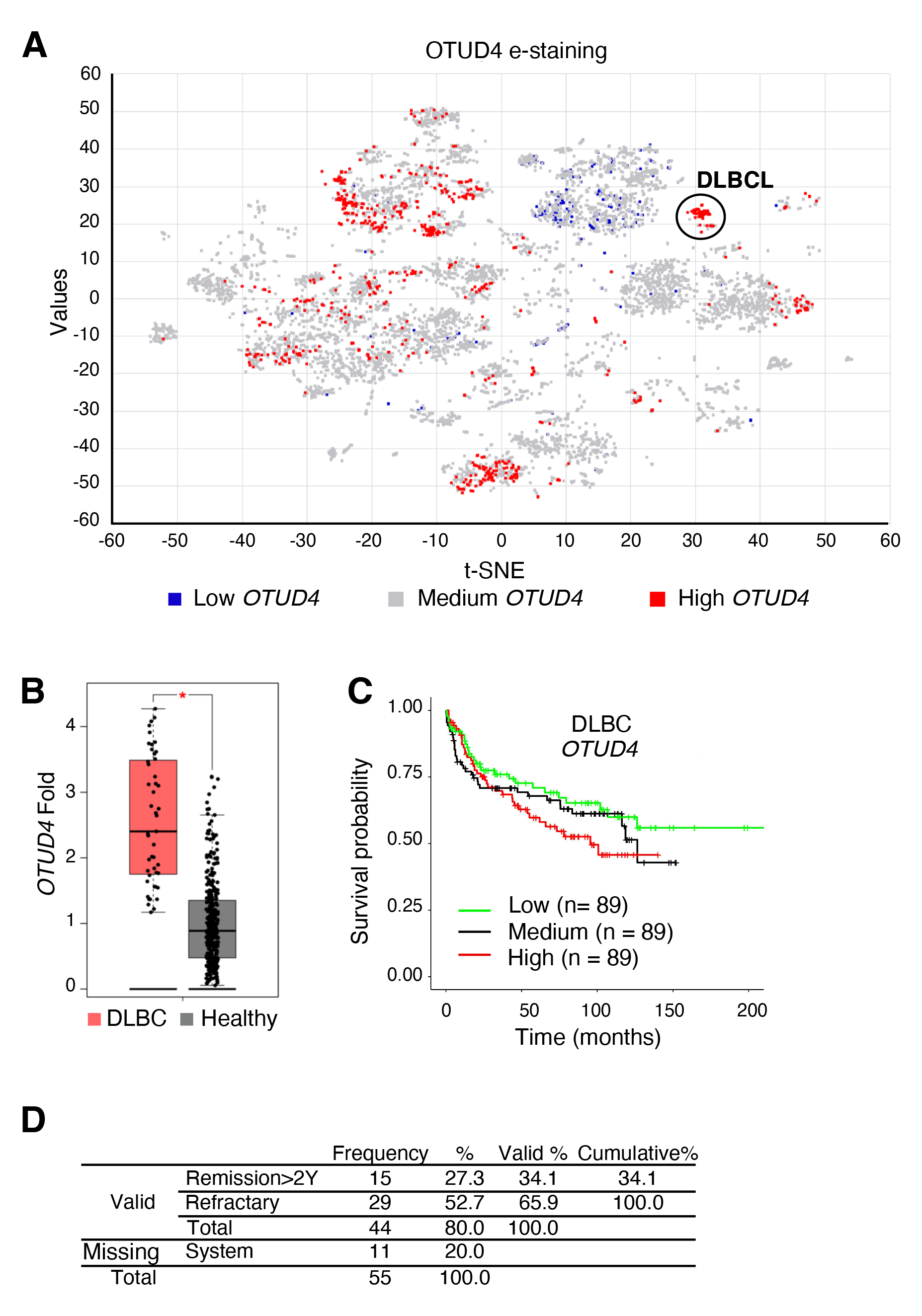
Gene expression of *OTUD4* in DLBCL. **A)** Significance of enrichment for *OTUD4* from different databases selected for visualization (e-staining) on hematologic cancer map using Hemap (http://hemap.uta.fi/). The *OTUD4* gene expression state is shown on the cancer map, where the color tones correspond to scaled log_2_ expression values (red, high; white, low; blue, not detected). DLBCL dataset is depicted inside a circle. **B)** Gene expression analysis of *OTUD4* in DLBC (Lymphoid Neoplasm Diffuse Large B cell Lymphoma) patients versus healthy volunteer group. p-value Cutoff 0,01. The box plot is blotted as a log_2_ (TPM+1) scale. Jitter Size 0,4. Data match TCGA normal and GTEx (gtexportal.org) data. Data retrieved from GEPIA pipeline (http://gepia.cancer-pku.cn). **C)** KM-plots of a cohort of 267 DLBCL patients. Cutoff-High and Low was 50%. Confidence interval 95%. **D)** Data corresponding to the DLBCL patient cohort (n=55) described in Figures 2E-F.

**Supplementary Fig. 5.**
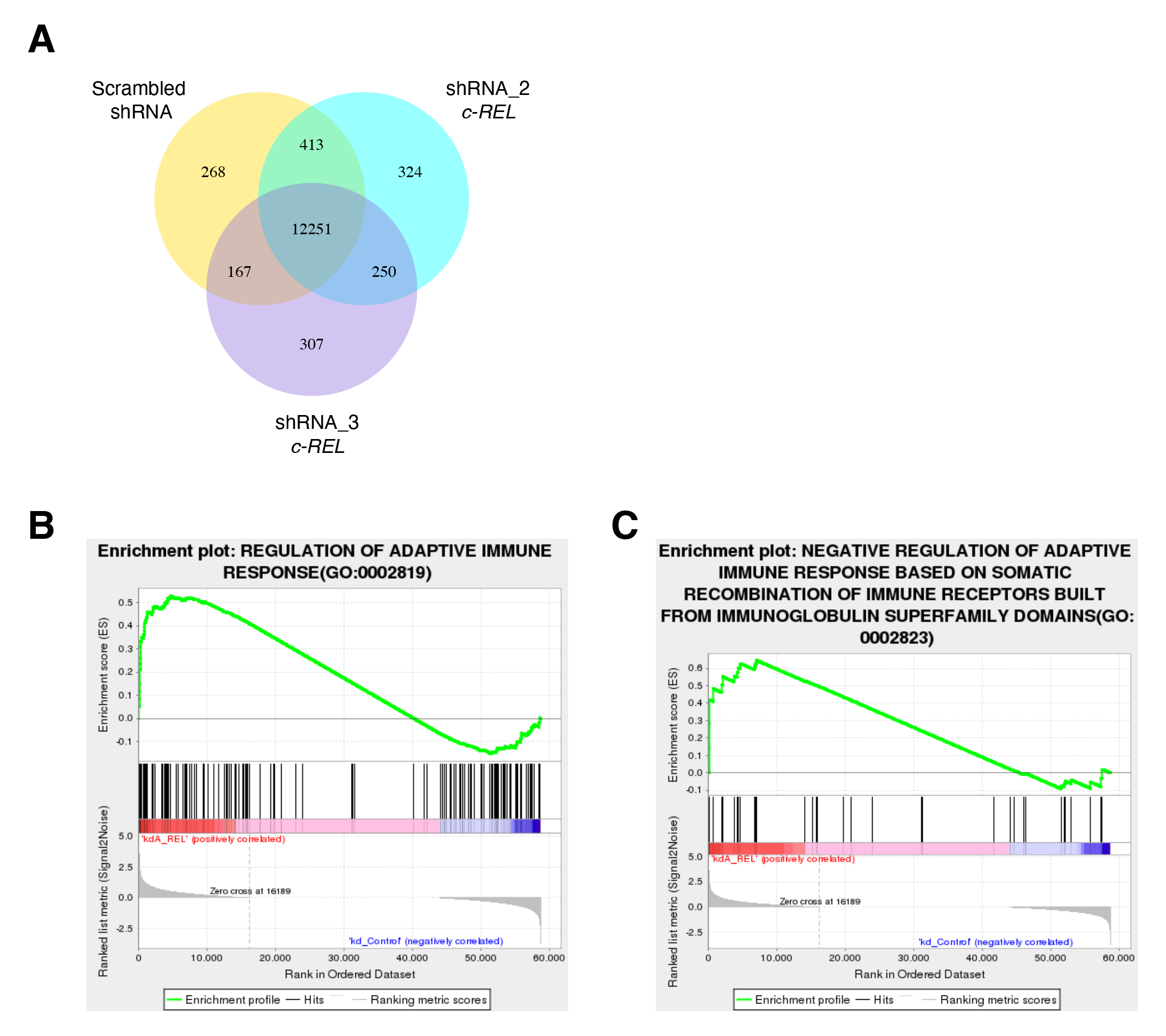
Enrichment analysis of the differential expressed genes in control and *c-REL* shRNA-treated BJAB cells. Coexpression Venn Diagram of RNA-Seq measurements. Numbers of genes uniquely expressed within Scrambled shRNA or two different shRNAs targeting *c-REL*, with the overlapping regions showing the number of genes that are co-expressed in two or more groups are depicted. See Supplementary Table 5. **B)** GSEA showing an enrichment of genes regulating *adaptive immune response* in BJAB cells upon *c-REL* downregulation when compared to control cells. Three independent experiments were subjected to RNA-seq analysis followed by GSEA. padj < 0.05 are considered significant enrichment. See Supplementary Table 6. **C)** Similar to B, GSEA showing an enrichment of genes in the *negative regulation of adaptive immune response based on somatic recombination of immune receptors built from immunoglobulin superfamily domains*. padj < 0.05 are considered significant enrichment. See Supplementary Table 6.

## Appendix. Supplementary Tables

Suppl. Table 1. Total proteome analysis in BJAB cells upon *c-REL* downregulation. Suppl. Table 2. RNA-seq analysis in BJAB cells upon *c-REL* downregulation.

Suppl. Table 3. Peaks of *OTUD4* found in ChIP-seq of c-REL in BJAB cells. Suppl. Table 4. *OTUD4* mRNA correlation to *c-REL* in blood cells.

Suppl. Table 5. Co-expression Venn Diagram. Control vs *c-REL* shRNAs in BJAB cells.

Suppl. Table 6. Enrichment analysis of the differential expressed genes upon *c-REL* downregulation in BJAB cells.

